# Influenza B viruses exhibit lower within-host diversity than influenza A viruses in human hosts

**DOI:** 10.1101/791038

**Authors:** Andrew L. Valesano, William J. Fitzsimmons, John T. McCrone, Joshua G. Petrie, Arnold S. Monto, Emily T. Martin, Adam S. Lauring

## Abstract

Influenza B virus undergoes seasonal antigenic drift more slowly than influenza A, but the reasons for this difference are unclear. While the evolutionary dynamics of influenza viruses play out globally, they are fundamentally driven by mutation, reassortment, drift, and selection within individual hosts. These processes have recently been described for influenza A virus, but little is known about the evolutionary dynamics of influenza B virus (IBV) at the level of individual infections and transmission events. Here we define the within-host evolutionary dynamics of influenza B virus by sequencing virus populations from naturally-infected individuals enrolled in a prospective, community-based cohort over 8176 person-seasons of observation. Through analysis of high depth-of-coverage sequencing data from samples from 91 individuals with influenza B, we find that influenza B virus accumulates lower genetic diversity than previously observed for influenza A virus during acute infections. Consistent with studies of influenza A viruses, the within-host evolution of influenza B viruses is characterized by purifying selection and the general absence of widespread positive selection of within-host variants. Analysis of shared genetic diversity across 15 sequence-validated transmission pairs suggests that IBV experiences a tight transmission bottleneck similar to that of influenza A virus. These patterns of local-scale evolution are consistent with influenza B virus’ slower global evolutionary rate.

**Importance:** The evolution of influenza virus is a significant public health problem and necessitates the annual evaluation of influenza vaccine formulation to keep pace with viral escape from herd immunity. Influenza B virus is a serious health concern for children, in particular, yet remains understudied compared to influenza A virus. Influenza B virus evolves more slowly than influenza A, but the factors underlying this are not completely understood. We studied how the within-host diversity of influenza B virus relates to its global evolution by sequencing viruses from a community-based cohort. We found that influenza B virus populations have lower within-host genetic diversity than influenza A virus and experience a tight genetic bottleneck during transmission. Our work provides insights into the varying dynamics of influenza viruses in human infection.

## Introduction

Influenza viruses rapidly mutate and evolve through selection, genetic drift, and reassortment (1). At a global scale, influenza A virus (IAV) and influenza B virus (IBV) evolve under strong positive selection driven by pressure for escape from pre-existing population immunity (2,3). Selection of new antigenic variants contributes to reduced effectiveness of seasonal influenza vaccines, necessitating annual updates of vaccine strains (4). IAV and IBV both undergo seasonal antigenic drift and share a similar genomic architecture, but their ecology and evolution differ in important ways (5). While IBV accounts for roughly one-third of influenza’s burden of morbidity and mortality (6,7), it circulates only in humans and is considered to be a lower pandemic risk than influenza A (IAV) due to the lack of an animal reservoir. Like IAV, there are co-circulating, antigenically-distinct lineages of IBV that are included in the quadrivalent influenza vaccine. Two lineages of IBV diverged in the 1980s, B/Victoria/2/87-like and B/Yamagata/16/88-like, here referred to as B/Victoria and B/Yamagata, respectively (8).

IBV evolves more slowly than IAV on a global scale and has a lower rate of antigenic drift, but the reasons for this are poorly understood (5,9). Similar evolutionary forces are involved in the antigenic evolution of both IAV and IBV, generally characterized by non-synonymous substitutions at antigenic sites in the surface hemagglutinin (HA) protein (10,11) and reassortment within and between lineages (12–14). The IBV polymerase has a lower mutation rate relative to IAV (15). However, it is unclear whether the slower global evolution of IBV is driven by its lower mutation rate or other differences in selection at the global scale.

All new seasonal influenza variants are ultimately derived from *de novo* mutations within individual hosts (16). Therefore, understanding how new variants arise within individuals and transmit between them is essential to defining how novel viruses spread in host populations. For example, if the relative mutation rate is a major factor underlying the global evolutionary differences across IAV and IBV, we might also expect to see differences in their within-host dynamics. We and others have used next-generation sequencing to investigate the within- and between-host evolutionary dynamics of IAV in humans (16–21). We have found that there is little accumulation of intrahost variants during acute infections of immunocompetent individuals (18,19), and we have not found evidence of changes in intrahost diversity by vaccination status or other proxies for immunological history (19,20,22). The IAV transmission bottleneck is stringent (19), which generally means that few variants that arise within hosts are able to transmit. Together, these studies suggest that selection of novel variants is an inefficient process in IAV-infected hosts, contrasting with its patterns of significant positive selection at the global level. Despite the importance of intrahost processes to influenza virus evolution, these dynamics have not been systematically investigated in IBV.

Here we use next-generation sequencing to define the within-host diversity of IBV populations from individuals enrolled in the Household Influenza Vaccine Evaluation (HIVE) study, a community-based household cohort initiated in 2010. We apply a previously-validated bioinformatic pipeline (23) to identify intrahost single-nucleotide variants (iSNV) arising during infection with B/Victoria and B/Yamagata viruses. We find that IBV has significantly lower intrahost diversity than IAV, consistent with its lower mutation rate and slower rate of evolution. We analyze shared iSNV across 15 genetically-validated household transmission pairs and find that, like IAV, IBV is also subject to a tight genetic bottleneck at transmission. These data provide the first systematic evaluation of the genetic architecture of IBV populations during natural human infection and provide insights into the comparative epidemiology and evolution of influenza viruses.

## Results

We used high depth-of-coverage sequencing to define the intrahost genetic diversity in IBV-positive samples collected from individuals in the HIVE, a prospective, household cohort in southeastern Michigan that follows 200-350 households annually (Table 1). This cohort provides an opportunity to investigate natural infections and transmission events in a community context. Individuals that meet symptom-based criteria for an upper respiratory illness during the surveillance period undergo collection of nasal and throat swabs for molecular detection of respiratory viruses by RT-PCR. Starting in 2014-2015, individuals also provided a sample collected at home prior to subsequent collection of a second specimen at the on-site clinic.

**Table 1.** Influenza B viruses over seven seasons in a household cohort.

|  | 2010-2011 | 2011-2012 | 2012-2013 | 2013-2014 | 2014-2015 | 2015-2016 | 2016-2017 |
| --- | --- | --- | --- | --- | --- | --- | --- |
| Households | 328 | 213 | 321 | 232 | 340 | 227 | 208 |
| Participants | 1441 | 943 | 1426 | 1049 | 1431 | 996 | 890 |
| Vaccinated n(%)<sup>a</sup> | 934 (65) | 554 (59) | 942 (66) | 722 (69) | 992 (69) | 681 (68) | 611 (69) |
| IBV Positive Individuals<sup>b</sup> | 45 | 7 | 49 | 4 | 44 | 11 | 30 |
| B/Yamagata | 1 | 3 | 38 | 4 | 34 | 5 | 26 |
| B/Victoria | 37 | 0 | 10 | 0 | 10 | 6 | 4 |
| IBV Positive Households<sup>c</sup> |  |  |  |  |  |  |  |
| Two Individuals | 10 | 2 | 5 | 0 | 11 | 2 | 4 |
| Three Individuals | 0 | 1 | 1 | 0 | 1 | 0 | 2 |
| High Quality NGS Data<sup>d</sup> | 13 | 2 | 20 | 1 | 32 | 11 | 20 |

<sup>a</sup> Self-reported or confirmed receipt of vaccine prior to the specified season.
<sup>b</sup> RT-PCR confirmed infection.
<sup>c</sup> Households in which two individuals were positive within 7 days of each other. In cases of trios, the putative chains could have no pair with onset >7 days apart.
<sup>d</sup> Samples with >10<sup>3</sup> genome copies per µl of transport medium, adequate amplification of all 8 genomic segments, and average sequencing coverage >10<sup>3</sup> per nucleotide.

Over seven seasons (2010-2011 through 2016-2017) and 8176 person-seasons of observation, we identified 111 individuals infected with B/Yamagata and 67 infected with B/Victoria (Table 1). Several households had clusters of infections of two or three IBV-positive individuals within 7 days of each other, suggestive of within-household transmission. Because variant identification is sensitive to input viral titer (23), we first measured viral loads of all available IBV-positive samples by RT-qPCR (Figure 1A). Any samples with a viral load below 10^3^ copies/μL were not submitted for sequencing. For samples with a viral load in the range of 10^3^-10^5^ copies/μL, we performed two independent RT-PCR reactions and sequenced replicate libraries on separate sequencing runs. We sequenced samples with viral loads above 10^5^ copies/μL of transport media in a single replicate. From the available IBV-positive samples, we were able to obtain sequence data on 106 samples from 91 individuals, consisting of 35 individuals infected with B/Victoria and 56 infected with B/Yamagata (Table 1).

**Figure 1.**
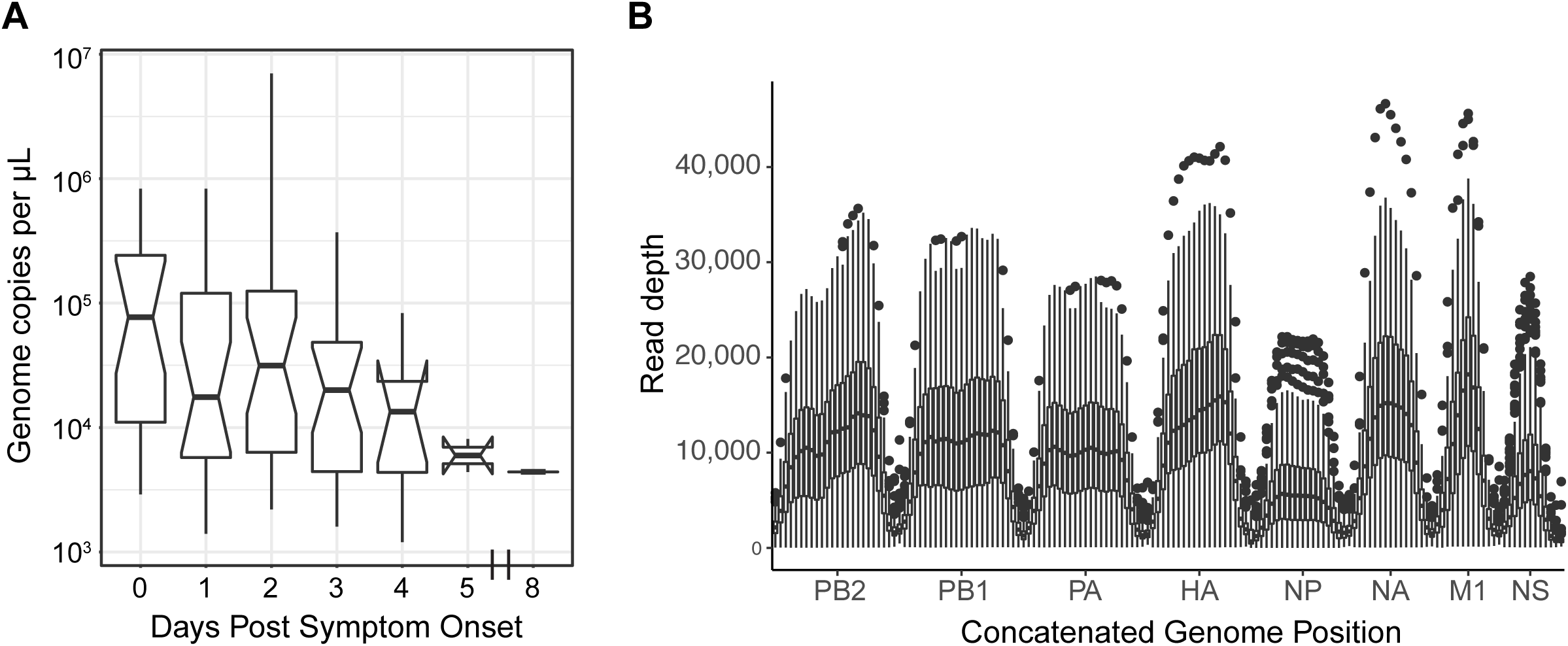
Viral load and sequencing coverage. (A) Boxplot of viral load (genome copies per microliter of swab transport media, y-axis) by day of sampling relative to symptom onset (x-axis). The boxes display median and 25^th^ and 75^th^ percentiles, with whiskers extending to the most extreme point within the range of the median ± 1.5 times the interquartile range; all values outside this range are shown as individual points. (B) Sequencing coverage is plotted with read depth on the y-axis and location within a concatenated influenza B virus genome on the x-axis. The mean coverage for each sample was calculated over a sliding window of size 200 and a step size of 100. The data are displayed for all samples at each window as a boxplot, showing the median and 25^th^ and 75^th^ percentiles, with whiskers extending to the most extreme point within the range of the median ± 1.5 times the interquartile range; all values outside this range are shown as individual points.

We identified intrahost single nucleotide variants (iSNV) using our previously-validated bioinformatic pipeline. As in our previous work, we report iSNV at frequencies of 2% or above, for which we have well-defined sensitivity and specificity (19). We consider sites with >98% frequency to be essentially fixed, setting the frequency at those sites to 100% (see Materials and Methods). We achieved a mean coverage of 10,000x per sample across most genome segments, with generally lower coverage on segments encoding NP and NS (Figure 1B). We restricted our analysis of iSNV to samples with an average genome coverage of greater than 1000x, which includes 99 of the original 106 sequenced samples.

### Within-host genetic diversity of IBV in natural infections

All samples exhibited low genetic diversity. The vast majority had no iSNV above the 2% cutoff. Of the 99 samples with high-quality NGS data, 70 had no minority iSNV, 17 had one iSNV, 7 had two iSNV, and 3 samples had 3 iSNV (median 0, IQR 0-2; Table 2). Two outliers had a large number of iSNV, with 8 and 20 iSNV. These two samples came from the same individual, with one collected at home and the second at the study clinic. Most of the iSNV in these two samples were present at similar frequencies, 3-5% and 17-23% in each specimen, respectively (Table 3), both of which were sequenced in duplicate on separate Illumina runs. The high number of mutations present at similar frequencies is suggestive of a mixed infection with distinct haplotypes or strains as opposed to *de novo* mutations arising on a single genetic background. The iSNV in the home-collected sample are all found in the subsequent clinic-collected sample, each with a similar change in frequency across the two samples. This further supports the conclusion that these mutations are on the same genome in a mixed infection with two distinct strains.

**Table 2.** Identified iSNV, excluding samples from one putative mixed infection.

| Enrollee | Specimen | Season | Lineage | Viral Load<sup>a</sup> | Gene | Nucleotide<sup>b</sup> | Amino Acid<sup>c</sup> | Frequency | Coverage<sup>d</sup> | Mutation Type<sup>e</sup> | Vaccinate<sup>f</sup> |
| --- | --- | --- | --- | --- | --- | --- | --- | --- | --- | --- | --- |
| 50207 | MH15919 | 16/17 | B/Victoria | 3.50E+03 | M | A650G | N207S | 0.024 | 13574 | NS | IIV4 |
| 331001 | MH2671 | 12/13 | B/Victoria | 3.20E+05 | NA | C276T | L73F | 0.086 | 6107 | NS | LAIV3 |
| 331001 | MH2671 | 12/13 | B/Victoria | 3.20E+05 | PA | A2047G | K671R | 0.474 | 5024 | NS | LAIV3 |
| 330171 | MH3227 | 12/13 | B/Victoria | 7.00E+06 | NA | A385C | N109T | 0.162 | 14964 | NS | IIV3 |
| 330171 | MH3227 | 12/13 | B/Victoria | 7.00E+06 | PA | C1982T | A649A | 0.461 | 8159 | S | IIV3 |
| 301587 | M53957 | 10/11 | B/Victoria | 3.30E+04 | HA | G1603A | G522R | 0.081 | 34517 | NS | No |
| 301587 | M53957 | 10/11 | B/Victoria | 3.30E+04 | NA | A863C | T268T | 0.038 | 45106 | S | No |
| 301202 | M54308 | 10/11 | B/Victoria | 4.40E+04 | PA | C1037T | N334N | 0.195 | 20857 | S | No |
| 50003 | MH10403 | 14/15 | B/Victoria | 8.20E+04 | NS | A103C | T18T | 0.063 | 3245 | S | No |
| 50004 | MH10404 | 14/15 | B/Victoria | 8.20E+04 | NP | A577G | N171S | 0.057 | 13413 | NS | No |
| 50004 | MH10404 | 14/15 | B/Victoria | 8.20E+04 | NA | A1457G | L466L | 0.223 | 4398 | S | No |
| 50004 | MH10404 | 14/15 | B/Victoria | 8.20E+04 | PA | G1617A | V528M | 0.497 | 17758 | NS | No |
| 50424 | HS1876 | 14/15 | B/Victoria | 1.60E+03 | NP | G1191A | D376N | 0.034 | 1073 | NS | IIV4 |
| 50051 | HS1909 | 14/15 | B/Victoria | 1.90E+03 | M | G709A | E227K | 0.343 | 2138 | NS | Yes, Unk |
| 50004 | HS1788 | 14/15 | B/Victoria | 8.30E+05 | NP | A577G | N171S | 0.054 | 12857 | NS | No |
| 50004 | HS1788 | 14/15 | B/Victoria | 8.30E+05 | PA | G1617A | V528M | 0.389 | 16217 | NS | No |
| 50004 | HS1788 | 14/15 | B/Victoria | 8.30E+05 | NA | A1457G | L466L | 0.045 | 3259 | S | No |
| 50312 | HS2019 | 15/16 | B/Victoria | 2.00E+05 | NP | G987A | V308I | 0.420 | 6593 | NS | No |
| 50312 | HS2019 | 15/16 | B/Victoria | 2.00E+05 | PA | G1346A | E437E | 0.467 | 9225 | S | No |
| 51123 | HS2680 | 15/16 | B/Victoria | 3.50E+05 | NP | G1511A | R482R | 0.344 | 7393 | S | No |
| 320779 | MH0776 | 11/12 | B/Yamagata | 3.40E+05 | NP | A735G | S223S | 0.023 | 9674 | S | IIV3 |
| 320779 | MH0776 | 11/12 | B/Yamagata | 3.40E+05 | PB2 | G661A | R211R | 0.222 | 14675 | S | IIV3 |
| 51092 | MH10076 | 14/15 | B/Yamagata | 1.20E+04 | PB1 | A223G | I66V | 0.116 | 7370 | NS | IIV4 |
| 50650 | MH16167 | 16/17 | B/Yamagata | 5.20E+04 | PA | T2019C | L662L | 0.373 | 5397 | S | IIV4 |
| 50650 | MH16167 | 16/17 | B/Yamagata | 5.20E+04 | PB1 | C345T | A106A | 0.159 | 6639 | S | IIV4 |
| 331060 | MH3065 | 12/13 | B/Yamagata | 3.70E+05 | PA | A1912G | K626R | 0.051 | 10393 | NS | LAIV3 |
| 331397 | MH4247 | 12/13 | B/Yamagata | 2.40E+04 | PB2 | A676G | R216R | 0.370 | 20110 | S | IIV3 |
| 330459 | MH4289 | 12/13 | B/Yamagata | 2.10E+05 | HA | G1102A | A355T | 0.024 | 14033 | NS | IIV3 |
| 330460 | MH4364 | 12/13 | B/Yamagata | 2.10E+05 | PB2 | G520A | V164V | 0.032 | 7919 | S | IIV3 |
| 50006 | MH16139 | 16/17 | B/Yamagata | 1.20E+05 | HA | T728C | F230S | 0.148 | 14877 | NS | No |
| 331471 | MH2216 | 12/13 | B/Yamagata | 8.60E+04 | PB2 | G1936A | Q636Q | 0.024 | 30012 | S | No |
| 331470 | MH2246 | 12/13 | B/Yamagata | 1.20E+04 | PA | G1535A | A500A | 0.029 | 9483 | S | No |
| 331470 | MH2246 | 12/13 | B/Yamagata | 1.20E+04 | PB2 | A2253G | K742R | 0.124 | 4478 | NS | No |
| 331470 | MH2246 | 12/13 | B/Yamagata | 1.20E+04 | PB2 | C769T | H247H | 0.023 | 12295 | S | No |
| 331364 | MH4166 | 12/13 | B/Yamagata | 2.80E+04 | HA | C746T | T236I | 0.037 | 29665 | NS | No |
| 331364 | MH4166 | 12/13 | B/Yamagata | 2.80E+04 | PA | G1298A | L421L | 0.093 | 15898 | S | No |
| UM41536 | MH6592 | 13/14 | B/Yamagata | 2.00E+04 | PB1 | G1893A | R622R | 0.022 | 20159 | S | No |
| 51093 | HS1747 | 14/15 | B/Yamagata | 3.90E+04 | PA | G1433A | L466L | 0.087 | 5646 | S | IIV4 |
| 50419 | HS3214 | 16/17 | B/Yamagata | 5.40E+05 | PB1 | C345T | A106A | 0.046 | 7545 | S | IIV4 |
| 51121 | HS3258 | 16/17 | B/Yamagata | 1.10E+04 | PB1 | A2079G | E684E | 0.022 | 17567 | S | No |

<sup>a</sup> Viral load measured by RT-qPCR, expressed in genome copies per microliter of transport medium.
<sup>b</sup> Consensus nucleotide followed by position on reference genome and variant nucleotide.
<sup>c</sup> Consensus amino acid followed by codon position on reference genome and variant amino acid.
<sup>d</sup> Coverage expressed as the total sequencing read depth at the site of the identified variant.
<sup>e</sup> Nonsynonymous (NS) or synonymous (S) variant relative to sample consensus.
<sup>f</sup> Self-reported or confirmed receipt of vaccine prior to the specified season. IIV4, quadrivalent inactivated; LAIV3, trivalent live attenuated; IIV3, trivalent inactivated; Unk, vaccine product unknown.

**Table 3.** Identified iSNV in one vaccinated individual^a^ with a putative mixed infection during the 2014-2015 season.

| Specimen<sup>a</sup> | Viral Load<sup>b</sup> | Gene | Nucleotide<sup>c</sup> | Amino Acid<sup>d</sup> | Frequency | Coverage<sup>e</sup> | Mutation Type<sup>f</sup> |
| --- | --- | --- | --- | --- | --- | --- | --- |
| HS1875 | 2.90E+04 | HA | G1061A | R341K | 0.029 | 22065 | NS |
| HS1875 | 2.90E+04 | NP | G1666A | G534D | 0.027 | 4688 | NS |
| HS1875 | 2.90E+04 | NA | G1210A | G384D | 0.054 | 20907 | NS |
| HS1875 | 2.90E+04 | NA | A798G | S247G | 0.052 | 25817 | NS |
| HS1875 | 2.90E+04 | NS | G1004A | V100V | 0.045 | 1944 | S |
| HS1875 | 2.90E+04 | PB2 | G817A | V263V | 0.036 | 24333 | S |
| HS1875 | 2.90E+04 | PB2 | A1231C | I401I | 0.032 | 26905 | S |
| HS1875 | 2.90E+04 | PB2 | A793G | E255E | 0.035 | 23840 | S |
| MH10536 | 3.80E+04 | HA | G1061A | R341K | 0.236 | 21341 | NS |
| MH10536 | 3.80E+04 | HA | C366T | C109C | 0.213 | 17059 | S |
| MH10536 | 3.80E+04 | M | A114G | L28L | 0.189 | 2792 | S |
| MH10536 | 3.80E+04 | NP | G1666A | G534D | 0.193 | 3039 | NS |
| MH10536 | 3.80E+04 | NP | G1257A | R398R | 0.166 | 11007 | S |
| MH10536 | 3.80E+04 | NA | G1210A | G384D | 0.239 | 20531 | NS |
| MH10536 | 3.80E+04 | NA | C1286T | Y409Y | 0.218 | 16887 | S |
| MH10536 | 3.80E+04 | NA | C1319T | C420C | 0.217 | 12891 | S |
| MH10536 | 3.80E+04 | NA | G1148A | R363R | 0.225 | 23087 | S |
| MH10536 | 3.80E+04 | NA | A798G | S247G | 0.233 | 24631 | NS |
| MH10536 | 3.80E+04 | NA | T816C | F253L | 0.06 | 24779 | NS |
| MH10536 | 3.80E+04 | NS | G1004A | V100V | 0.185 | 1310 | S |
| MH10536 | 3.80E+04 | NS | G596A | V183I | 0.190 | 10074 | NS |
| MH10536 | 3.80E+04 | NS | T469A | V140V | 0.198 | 8807 | S |
| MH10536 | 3.80E+04 | NS | T66C | M6T | 0.173 | 1986 | NS |
| MH10536 | 3.80E+04 | PA | G1279A | S415N | 0.188 | 19813 | NS |
| MH10536 | 3.80E+04 | PB1 | T1932A | S635S | 0.214 | 19999 | S |
| MH10536 | 3.80E+04 | PB2 | G817A | V263V | 0.217 | 17558 | S |
| MH10536 | 3.80E+04 | PB2 | A1231C | I401I | 0.218 | 19869 | S |
| MH10536 | 3.80E+04 | PB2 | A793G | E255E | 0.221 | 17325 | S |

<sup>a</sup> Enrollee number 50425. HS Indicates home specimen and MH indicates clinic specimen, both from same individual
<sup>b</sup> Viral load measured by RT-qPCR, expressed in genome copies per microliter of transport medium.
<sup>c</sup> Consensus nucleotide followed by position on reference genome and variant nucleotide.
<sup>d</sup> Consensus amino acid followed by codon position on reference genome and variant amino acid.
<sup>e</sup> Coverage expressed as the total sequencing read depth at the site of the identified variant.
<sup>f</sup> Nonsynonymous (NS) or synonymous (S) variant relative to sample consensus.

We examined how within-host diversity changes by day of sampling during IBV infections, as the virus population rapidly expands and contracts. As we have previously shown that specimen viral load can affect the sensitivity and specificity of variant identification (23), we sought to control for this variable in our analysis. Although viral load generally decreased with time after symptom onset (Figure 1A), we found that within-host diversity as measured by number of identified minority iSNV did not vary with viral load (Figure 2A) or with day of infection (Figure 2B). The frequencies of the identified iSNV were consistent across replicate libraries from the same samples, indicating that our measurements of iSNV frequency are precise (Figure 2C).

**Figure 2.**
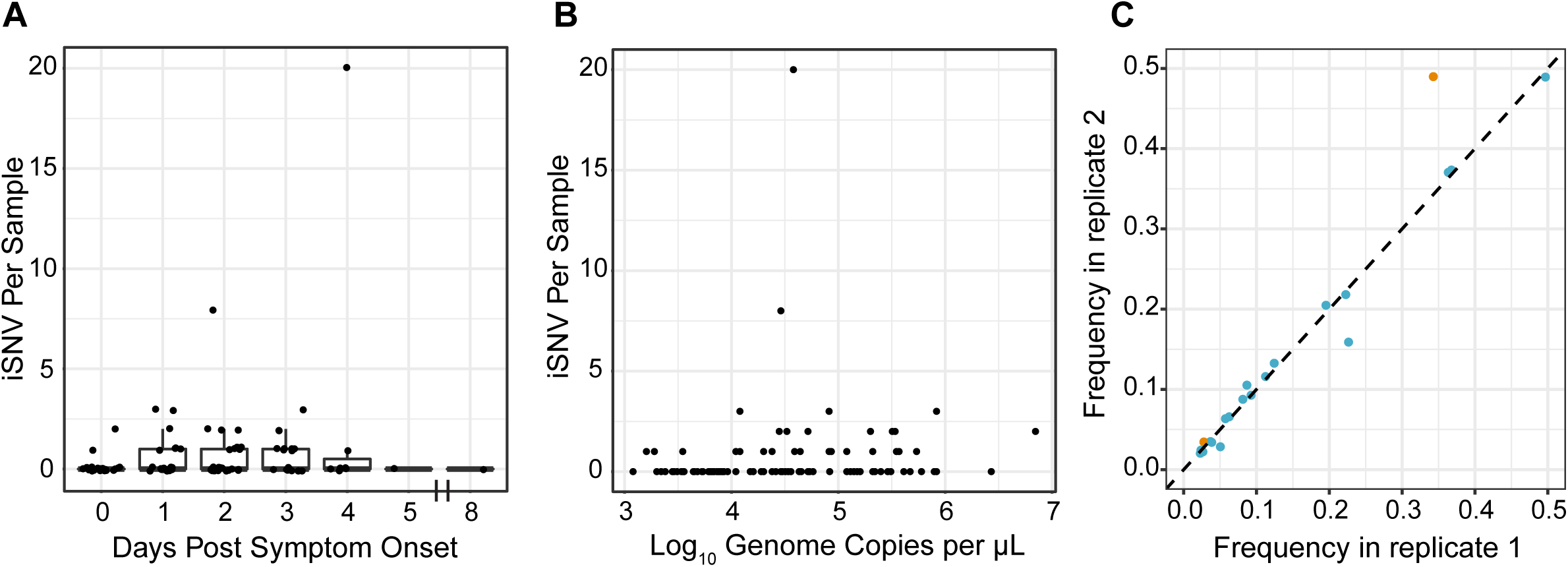
Intrahost minority SNV by day post-symptom onset and viral load. (A) Number of minority iSNV per sample is plotted on the y-axis by day post symptom onset on the x-axis. Data are displayed as boxplots representing the median and 25^th^ and 75^th^ percentiles, with whiskers extending to the most extreme point within the range of the median ±1.5 times the interquartile range. The raw data points are overlaid on top of the boxplots. (B) Scatterplot relating the number of minority iSNV per sample on the y-axis to the log_10_ of viral load, in genome copies per microliter, on the x-axis. (C) Frequency of minority iSNV in samples sequenced in duplicate. Orange dots represent variants identified in samples with viral load of 10^3^ – 10^4^ genome copies per microliter and blue dots represent variants in samples with viral load of 10^4^ – 10^5^ genome copies per microliter.

We detected minority iSNV across all eight genome segments (Figure 3). We identified more synonymous than nonsynonymous mutations, which given the ratio of synonymous and nonsynonymous sites indicates that purifying selection dominates within hosts. There was only one minority iSNV present in more than one individual; we identified a variant encoding a synonymous mutation in PB1 in two individuals from separate households infected with B/Yamagata in the 2016-2017 season. We did not identify any nonsynonymous minority iSNV in the known antigenic sites of IBV hemagglutinin, which suggests that positive selective pressure for variants that escape antibody-mediated immunity is not particularly strong within hosts. We found that there is no difference in the distribution of the number of iSNV per sample between vaccinated and non-vaccinated individuals (Figure 4A). During the first few seasons of the study, some individuals received trivalent vaccines, which contain only one of the two IBV lineages. We therefore repeated this analysis, excluding 3 individuals for whom we had no information about specific vaccine product and re-classifying 6 individuals who received trivalent vaccines and were infected with a lineage not included in that season’s trivalent formulation as “unvaccinated.” We again found no difference in the number of iSNV between groups (MWU test, p = 0.9103). Together, these data indicate that vaccine-induced immunity is not a major diversifying force for IBV within hosts in our study population. This is consistent with our previous work on IAV in the HIVE as well as a randomized-controlled trial of vaccine efficacy (FLU-VACS), both of which showed no difference in intrahost diversity based on same-season vaccination status (19,20). Intrahost diversity was similar between B/Victoria and B/Yamagata virus populations (Figure 4B), consistent with our previous comparison of subtype A/H3N2 and A/H1N1 viruses (19).

**Figure 3.**
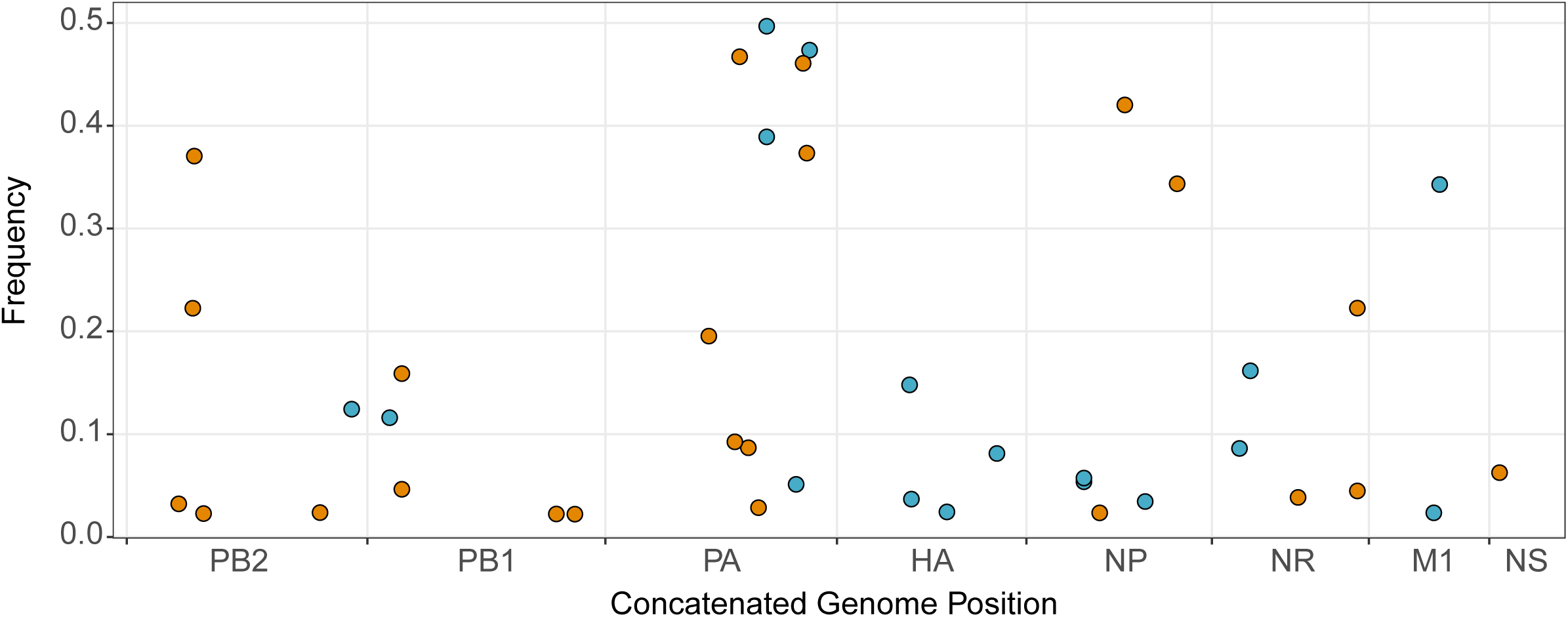
Intrahost SNV frequency by genome position and mutation type. All minority (<50%) iSNV from 99 samples are displayed with their frequency on the y-axis and their position within a concatenated influenza B virus genome on the x-axis. Synonymous mutations are shown in orange and nonsynonymous mutations in blue.

**Figure 4.**
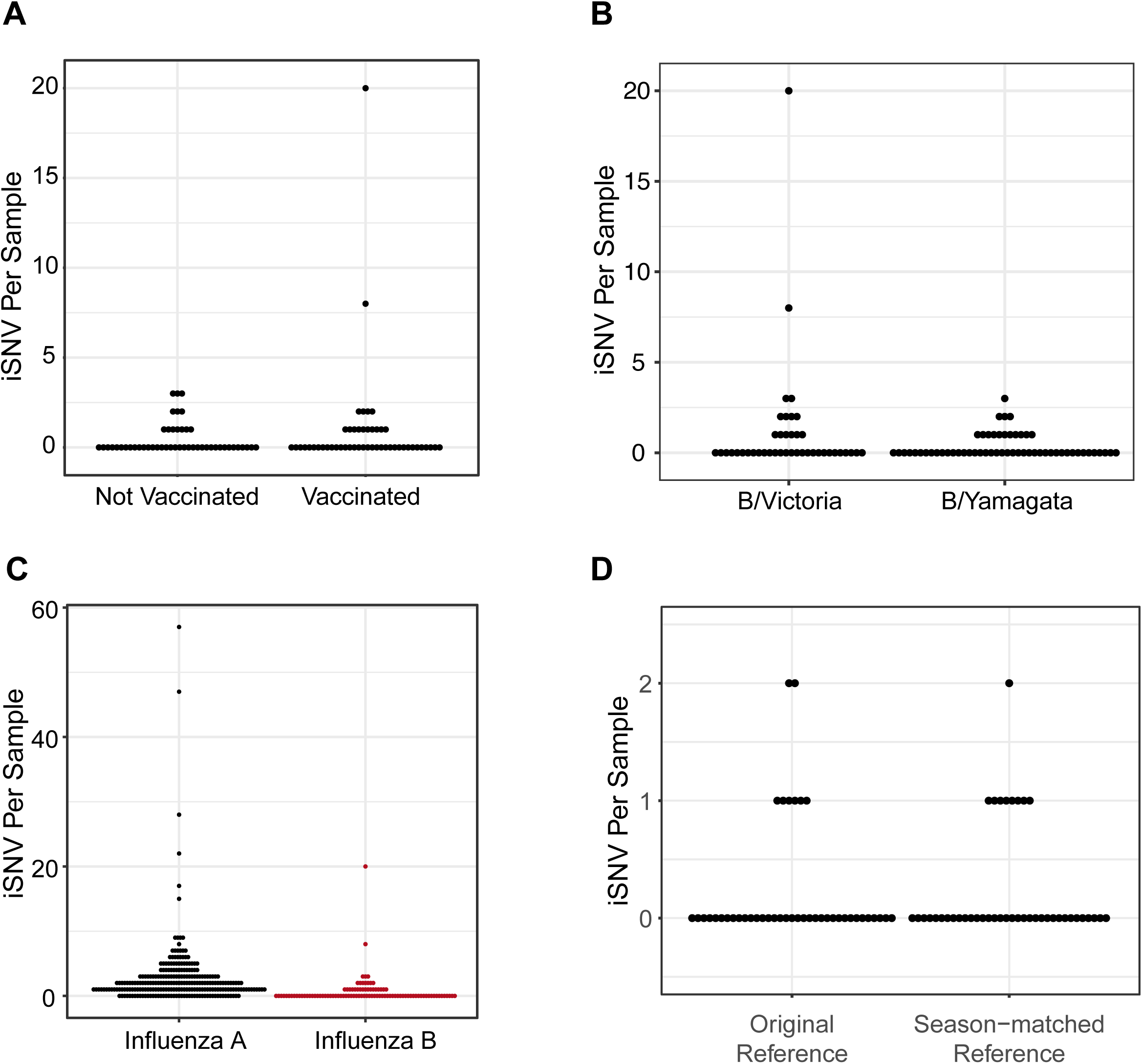
Intrahost SNV by vaccination status and IBV lineage. (A) Numbers of minority iSNV per sample across all 99 samples are shown (y-axis) by current-season vaccination status of the host (x-axis). (B) Numbers of minority iSNV per sample are shown (y-axis) by IBV lineage (x-axis). (C) Numbers of minority iSNV per sample are shown (y-axis) by influenza virus type (x-axis). Data for influenza A virus are from 249 samples described in McCrone et al. 2018. Data on influenza B virus are from 99 high-quality samples in the present study. (D) Numbers of minority iSNV in 43 of the 99 high-quality samples (y-axis), consisting of B/Yamagata from the 2014/2015 season, B/Victoria from the 2015/2016 season, and B/Yamagata from the 2016/2017 season based on alignments to the original references from the 2012/2013 season vs. season-matched reference genomes (x-axis).

We compared the within-host genetic diversity of IBV to our previously published data on IAV from the HIVE cohort (19). Here, IBV exhibits lower within-host diversity compared to IAV (Figure 4C). To ensure our results were not an artifact of overly stringent quality thresholds, we also identified minority iSNV with less conservative read mapping quality (MapQ) and base quality (Phred) scores. We identified the same set of minority iSNV with a MapQ cut-off of 20 as with the original cutoff of 30. Similarly, reduction of the Phred base-quality cutoff to >25 in addition to a MapQ score cutoff of >20 resulted in only 20 more minority iSNV, eight of which were found in the individual with a mixed infection. The other additional 12 minority iSNV were dispersed across specimens and did not significantly change the overall distribution of within-host diversity. We also examined whether our results were biased by use of a single B/Yamagata and B/Victoria reference for alignment and variant calling, which were both drawn from the 2012-2013 season (see Materials and Methods). We realigned sequence data from 43 of the original 99 samples to season-specific reference genomes isolated in southeastern Michigan. We found that the overall alignment rate for any given specimen was similar between the original reference and the new season-matched reference. Variant identification based on the new references and the original quality thresholds resulted in the same distribution of within-host diversity, although the identity of some iSNV was different (Figure 4D).

Together, these results indicate that our measurements of within-host diversity are robust to several technical aspects of variant identification and are unlikely to account for the lower observed diversity of IBV. Because these data are from the same cohort and were generated using the same sequencing approach and analytic pipeline as our previous IAV datasets, the observed differences likely reflect true biological differences between IAV and IBV.

### Identification of household transmission pairs

We compared viral diversity across samples from individuals in the same household to investigate the genetic bottleneck that influenza B viruses experience during natural transmission. Over the seven influenza seasons, thirty-nine households in the HIVE cohort had two or more individuals positive for the same IBV lineage within a 7-day interval (Table 1). This epidemiologic linkage is suggestive of transmission events but does not rule out co-incident community acquired infection (19). We identified 16 putative transmission pairs for which we sequenced at least one sample from each individual. In one of these pairs, the putative recipient was the individual with a mixed infection. The donor did not have evidence of a mixed infection based on number of iSNV, which would imply that the recipient may have been infected twice or that the second virus was lost from the donor by the time of sampling. This pair was excluded from the between-host analysis, leaving 15 putative transmission pairs for which we have high-quality sequencing data on both donor and recipient influenza populations.

We used our sequencing data to determine which of these epidemiologically linked household pairs were actual IBV transmission pairs. We generated maximum likelihood phylogenetic trees for samples from the two IBV lineages using the concatenated coding consensus sequences. Phylogenetic analysis provided genetic evidence that the 15 epidemiologically-linked pairs were indeed true transmission pairs, as epidemiologically-linked pairs were found nearest each other in each tree (Figure 5A and 5B; vertical bars with household ID). We also validated these transmission pairs by analyzing the genetic distance across viral populations. True transmission pairs should have genetically similar populations exhibiting low genetic distance, while individuals with coincident community acquisition are more likely to have populations with a higher genetic distance. We compared the genetic distance between epidemiologically-linked household pairs and random community pairs from the same season and infected with the same IBV lineage, using L1-norm as measurement of genetic distance (Figure 5C). The distribution of random community pairs functions as a null model of genetic distances among locally circulating strains. All of the 15 putative transmission pairs fell on the tail of this distribution, below the 5^th^ percentile of the community pair L1-norm distribution, indicating that they are true transmission pairs (Figure 5C). While the L1-norm is a function of both the consensus sequence and the iSNV, this signal was predominantly driven by consensus differences, as reflected in the phylogenetic analysis.

**Figure 5.**
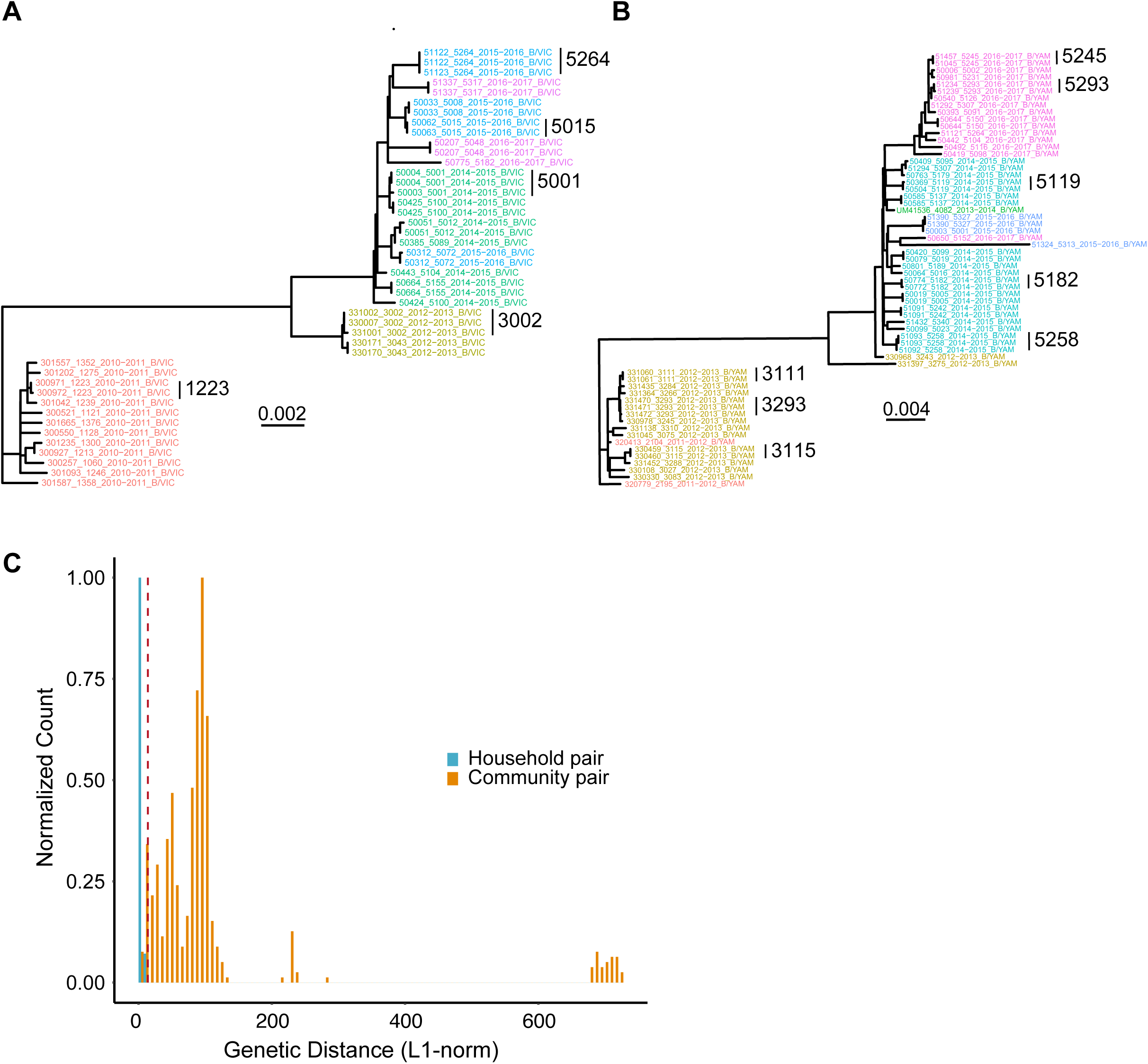
Identification of household transmission pairs. Maximum likelihood phylogenetic tree of all B/Victoria (A) and B/Yamagata (B) samples from this study. Concatenated consensus coding sequences were aligned with MUSCLE and phylogenetic trees constructed with RAxML. Tip labels are denoted as enrollee ID, household ID, season, and lineage, separated by underscores; tip labels are color-coded by household ID. (C) Histogram of genetic distance, as measured by L1-norm, between household pairs and random community pairs from the same season and lineage. The bar heights for each group are normalized to the maximum for each group for comparison. Community pairs are shown in orange and household pairs shown in blue. The dotted red line indicates the 5^th^ percentile of the community pair distribution.

### Comparison of viral diversity across transmission pairs

Transmission bottlenecks restrict the genetic diversity that is passed between hosts. With a loose transmission bottleneck, many unique genomes will be passed from donor to recipient. Because this will allow two variants at a given site to be transmitted, sites that are polymorphic in the donor are more likely to be polymorphic in the recipient. However, in the case of a tight or stringent bottleneck, sites that are polymorphic in the donor will likely be either fixed or absent in the recipient. We have previously demonstrated that influenza A experiences a tight transmission bottleneck of 1-2 unique genomes (19). Across our 15 IBV transmission pairs, we found no sites that were polymorphic in the donor and recipient (Figure 6). Intrahost SNV present in the donor were either fixed (100%) or absent (0%) in the recipient. These data suggest a stringent transmission bottleneck for influenza B, similar to that of influenza A. As there were fewer samples, transmission pairs, and iSNV in our IBV dataset, we were unable to obtain a robust and precise estimate of bottleneck size.

**Figure 6.**
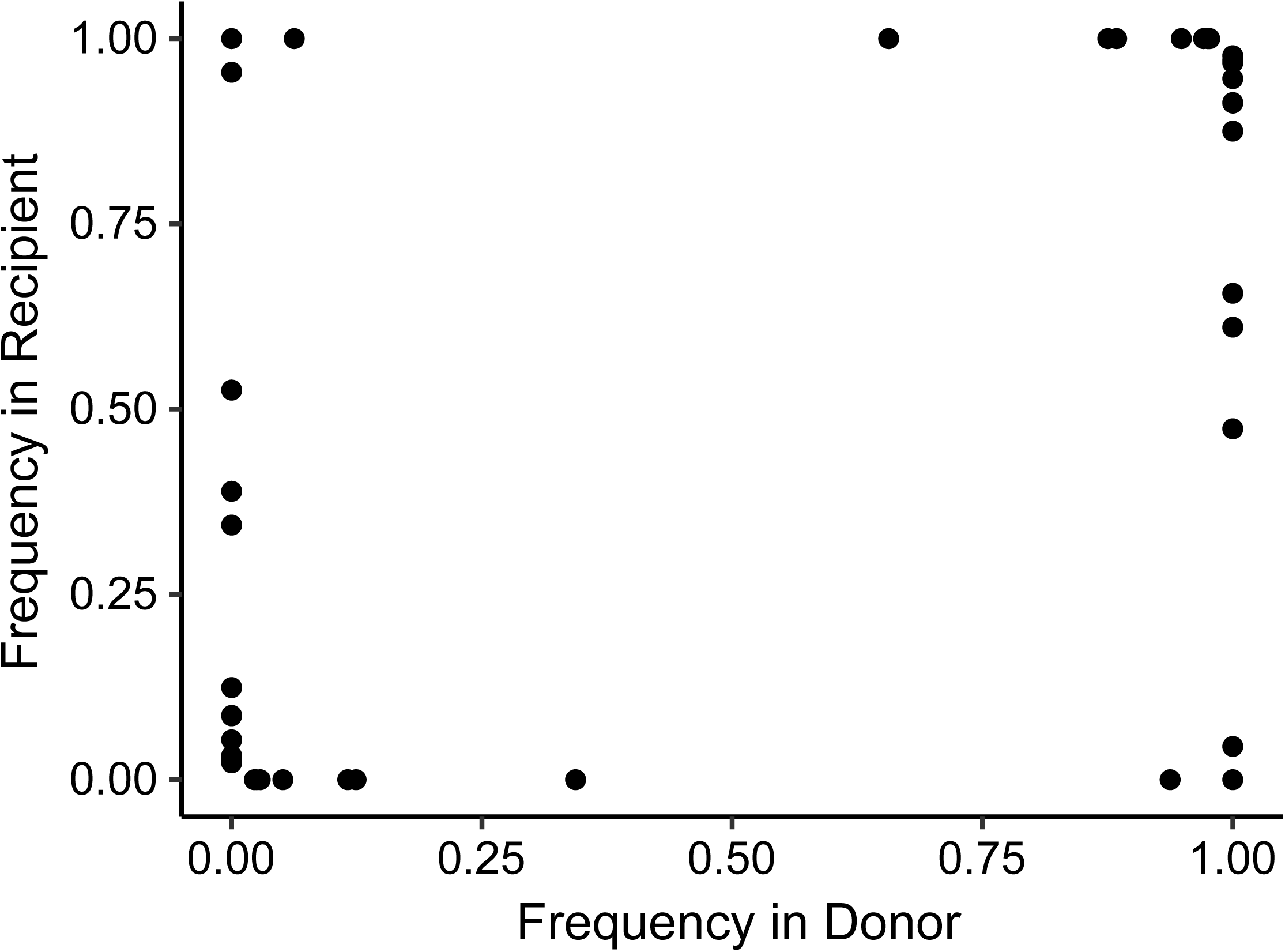
Shared diversity across household transmission pairs with influenza B virus. Intrahost SNV for 15 validated transmission pairs using samples closest to the time of transmission (inferred based on day of symptom onset). Each iSNV is plotted as a point with its frequency in the recipient (y-axis) versus its frequency in the donor (x-axis).

## Discussion

Here we define the within-host genetic diversity of IBV in natural infections by sequencing 106 samples collected over 8176 person-seasons of observation in a household cohort. Because the HIVE study prospectively identifies individuals with acute respiratory illness regardless of severity, these samples capture IBV dynamics in a natural setting, reflective of infections occurring in the community. We show that within-host diversity of IBV is remarkably low, with most samples displaying no intrahost variants above our level of detection. We also find that IBV experiences a tight transmission bottleneck, limiting the diversity that is passed between hosts. IBV exhibits significantly lower within-host diversity compared to IAV. These findings reflect the slower relative evolutionary rate of IBV compared to IAV.

Our findings are largely consistent with what has been observed in IAV infections in humans (17–20). We found that only a minority of samples contain iSNV, the majority of which encode synonymous changes, consistent with a predominance of purifying selection within hosts. If immune-driven selective pressures were sufficiently strong to drive positive selection of antigenic variants at the individual level, we would expect to see enrichment of variants in antigenic regions. However, variants were no more common in the antigenic proteins, hemagglutinin and neuraminidase, and we found no intrahost variants in known antigenic regions of hemagglutinin. We also found that the extent of within-host diversity did not vary with current-season vaccination status, further suggesting that immune selection is not particularly strong within hosts (19,20,22). Our data suggest that selective sweeps occur infrequently at the individual level, with selection only evident over a broader scale of time and space (19,35). We recognize, however, that it is possible for individual level selective pressure to vary in magnitude by age, locale, influenza infection history, or immune status (36).

We do find that there are important differences in the within-host evolution of IAV and IBV. IBV displays significantly lower within-host diversity compared to IAV. Since measurements of within-host diversity can vary based on host population, sequencing approach, and variant calling algorithm (37), a strength of our study is that our comparison is based on samples from the same cohort with the same sequencing approach and analytic pipeline. In both of our studies, we have sequenced swab samples directly without prior culture, accounted for the confounding effect of viral load, and used a standardized, empirically-validated analytic pipeline for variant identification (23). This pipeline includes rigorous quality criteria to reduce false positives that can be introduced by amplification and Illumina sequencing. Importantly, these empirical quality criteria did not mask diversity actually present in these samples, strengthening the conclusion that IBV exhibits lower within-host diversity compared to IAV.

The most likely biological explanation for IBV’s lower within-host diversity is its *de novo* mutation rate, which is thought to be at least two-fold lower than that of IAV (15). Viral mutation rates are critical to the diversification of rapidly evolving viruses within hosts. Under a neutral model, the number and frequency of minority variants is dependent on the mutation rate and demographics of the population (16). In such a model, the expected number of variants is highly sensitive to variation in the mutation rate across the range commonly estimated in RNA viruses. In light of our results, a more thorough comparison of mutation rates across influenza viruses is needed.

Another possible factor underlying IBV’s reduced diversity is the mutational robustness of the IBV genome relative to IAV. If IBV were less robust to mutation, stronger negative selection on multiple genes in IBV could result in more limited within-host diversity, perhaps located to certain regions of the genome. However, we found that the distributions of iSNV across IAV and IBV genomes are relatively similar. Furthermore, we have previously shown that the distribution of mutational fitness effects in influenza A/WSN/33/H1N1 matches that of other RNA and ssDNA viruses (38). Given that viruses across families with vastly different genomic architecture have similar mutational robustness, this is unlikely to account for the differences in within-host diversity between IAV and IBV.

We find that IBV experiences a stringent genetic bottleneck between hosts. A stringent transmission bottleneck places a constraint on the rate of adaptation of viral populations within and between individual hosts. Population bottlenecks reduce the effective population size, which increases random genetic drift and decreases the efficiency of selection (39). This results in a reduced ability of selection to fix beneficial mutations and to remove deleterious ones, which can decrease population fitness. However, there are potential evolutionary advantages to stringent bottlenecks, including removal of defective interfering particles (40,41). While we were not able to estimate the size of the transmission bottleneck as precisely as IAV, it is likely that the bottleneck size is comparable across the two viruses given the similarities in their transmission routes and ecology in the human population. Data from many more transmission pairs will be necessary for a more robust estimate.

Together, our results are consistent with the slower rate of global evolution observed in IBV lineages compared with both seasonal A/H1N1 and A/H3N2 (10,12,14,42). We suggest that a lower intrinsic mutation rate leads to reduced within-host diversity. With a comparably tight bottleneck, fewer *de novo* variants will rise to a level where they can be transmitted and spread through host populations. Combined with a lower incidence of IBV versus IAV, this would result in fewer variants that eventually spread and influence global dynamics. However, further investigation in larger populations will be required to evaluate the within-host dynamics of both types of seasonal influenza viruses and how they contribute to larger-scale evolutionary patterns.

## Materials and Methods

### Description of the HIVE cohort

The HIVE study is a prospective, community-based household cohort in Southeastern Michigan based at the University of Michigan School of Public Health (24–29). The cohort was initiated in 2010, with enrollment of households with children occurring on an annual basis and an active surveillance period lasting from October through May. In 2014, active surveillance was expanded to take place year-round. Participating adults provided informed consent for themselves and their children, and children ages 7-17 provided oral assent. Individuals in each household were followed prospectively for acute respiratory illness, defined as two or more of the following: cough, fever or feverishness, nasal congestion, chills, headache, body aches, or sore throat. Study participants meeting the criteria for acute respiratory illness attended a study research clinic at the University of Michigan School of Public Health where a combined throat and nasal swab, or a nasal swab only for children less than three years old, was collected by the study team. Beginning in the 2014-2015 season, study participants with acute respiratory illnesses took an additional nasal swab at home at the time of illness onset, collected either by themselves or by a parent. The study was approved by the Institutional Review Board of the University of Michigan Medical School.

### Viral detection, lineage typing, and viral load quantification

We processed upper respiratory specimens (combined nasal and throat swab or nasal swab) for confirmation of influenza virus infection by reverse transcription polymerase chain reaction (RT-PCR). We extracted viral RNA with either QIAamp Viral RNA Mini Kits (Qiagen) or PureLink Pro 96 Viral RNA/DNA Purification kits (Invitrogen) and tested samples using the SuperScript III Platinum One-Step Quantitative RT-PCR System with ROX (Invitrogen) and primers and probes for universal detection of influenza A and B (CDC protocol, 28 April 2009). Specimens positive for influenza virus were tested using subtype/lineage primer and probe sets, which are designed to detect influenza A (H3N2), A (H1N1)pdm09, B (Yamagata), and B (Victoria). An RNAseP primer/probe set was run for each specimen to confirm specimen quality and successful RNA extraction.

We quantified the viral load in each sample by RT-qPCR using primers specific for the open reading frame of segment 8 (NS1/NEP): forward primer 5’-TCCTCAACTCACTCTTCGAGCG-3’, reverse primer 5’-CGGTGCTCTTGACCAAATTGG-3’, and probe 5’-(FAM)- CCAATTCGAGCAGCTGAAACTGCGGTG-(BHQ1)-3’. Each reaction contained 5.4 μL of nuclease-free water, 0.5 μL of each primer at 50 μM, 0.1 μL of ROX dye, 0.5 μL SuperScript III RT/Platinum Taq enzyme mix, 0.5 μL of 10 μM probe, 12.5 μL of 2x PCR buffer master mix, and 5 μL of extracted viral RNA. To relate genome copy number to Ct value, we used a standard curve based on serial dilutions of a plasmid control, run in duplicate on the same plate.

### Amplification, library preparation, and sequencing

We amplified viral cDNA from all eight genomic segments using the SuperScript III One-Step RT-PCR Platinum Taq HiFi Kit (Invitrogen). Each reaction contained 5 μL of extracted viral RNA, 12.5 μL of 2x PCR buffer, 2 μL of primer cocktail, 0.5 μL of enzyme mix, 5 μL of nuclease-free water. The primer cocktail was a mixture of B-PBs-UniF, B-PBs-UniR, B-PA-UniF, B-PA-UniR, B-HANA-UniF, B-HANA-UniR, B-NP-UniF, B-NP-UniR, B-M-Uni3F, B-Mg-Uni3F, B-M-Uni3R, B-NS-Uni3F, and B-NS-Uni3R (sequences and proportions are listed in ref. (30)). The thermocycler protocol was: 45 °C for 60 min, 55 °C for 30 min, 94 °C for 2 min, then 5 cycles of 94 °C for 20 s, 40 °C for 30 s, 68 °C for 3 min 30 s, then 40 cycles of 94 °C for 20 s, 58 °C for 30 s, 68 °C for 3 min 30 s, and a final extension of 68 °C for 10 min. We confirmed IBV genome amplification by gel electrophoresis. We sheared amplified cDNA (100-500 ng) on a Covaris ultrasonicator with the following settings: time 80 sec, duty cycle 10%, intensity 4, cycles per burst 200. We prepared sequencing libraries with NEBNext Ultra DNA Library Prep kits (NEB) and sequenced them on an Illumina NextSeq with 2×150 paired end reads (mid-output run, v2 chemistry). To increase the specificity of variant identification, samples with a viral load between 10^3^ and 10^5^ genome copies/μL of transport media were amplified and sequenced in duplicate. Samples amplified from B/Victoria and B/Yamagata plasmid clones were included on each sequencing run to account for sequencing errors. The plasmids used in the control reactions were generated by segment-specific RT-PCR from clinical samples of B/Victoria and B/Yamagata strains from the 2012-2013 season followed by gel extraction and TOPO-TA cloning (Invitrogen). The sequence of each plasmid was determined by Sanger sequencing. We generated the plasmid control amplicons included on each Illumina sequencing run using the same multiplex amplification protocol, but with cloned plasmid DNA as the template.

### Identification of iSNV

Intrahost single-nucleotide variants (iSNV) were identified using a previously-described analytic pipeline (23). We identified iSNV in samples that had an average genome coverage greater than 1000x and a viral load greater than 10^3^ genome copies per microliter of transport media in the original sample. Sequencing adapters were removed with cutadapt (31) and reads were aligned to the sequences derived from the B/Victoria and B/Yamagata plasmid controls with Bowtie2 (32). Duplicate reads were marked and removed with Picard and samtools (33). Putative variants were identified with the R package deepSNV using data from the clonal plasmid controls of each sequencing run (34). Minority iSNV (<50% frequency) were identified using the following empirically-derived criteria: deepSNV p-value <0.01, average mapping quality >30, average Phred score >35, and average read position in the middle 50% (positions 37 and 113 for 150 base pair reads). For samples processed in duplicate, we used only variants that were present in both replicates; the frequency of the variant in the replicate with greater coverage at that site was used. Lastly, variants with frequency <2%, which have a higher false positive rate from RT-PCR and/or sequencing errors, were not included in downstream analyses.

In our previous work on IAV, we found that there were multiple sites with mutations that were essentially fixed (>0.95) relative to the plasmid control and in which the base in the plasmid control was therefore identified as a minority variant in the sample (19). At these sites, deepSNV is unable to estimate the base-specific error rate and cannot distinguish true minority iSNV; however, we found that we could accurately identify minority variants at these sites at a frequency of 2% or above (19). This frequency threshold was incorporated into the pipeline for iSNV identification at these sites. Therefore, we report intrahost variants from 2-98%; minority iSNV are the subset of these variants with a frequency between 2-50%. Any sites that were monomorphic after applying quality filters were assigned a frequency of 100%.

### Data and code availability

Raw sequence data, with human content filtered out, are available at the NCBI Sequence Read Archive under BioProject accession number PRJNA561158. Code for the variant identification pipeline is available at http://github.com/lauringlab/variant_pipeline. Analysis code is available at http://github.com/lauringlab/Host_level_IBV_evolution.

## Acknowledgements

We acknowledge the individuals enrolled in the HIVE study for their participation. We thank Maria Virgilio for technical assistance. This work was supported by NIH R01 AI 118886, NIH R21 AI 141832 and a Burroughs Wellcome Fund Investigator in the Pathogenesis of Infectious Diseases award to ASL. The HIVE cohort was supported by CDC U01 IP 001034 to ASM and ETM, and CDC U01 IP000474 and NIH R01 097150 to ASM. ALV was supported by NIH T32 GM 007863.

